# ODAMNet: a Python package to identify molecular relationships between chemicals and rare diseases using overlap, active module and random walk approaches

**DOI:** 10.1101/2023.07.05.546536

**Authors:** Morgane Térézol, Anaïs Baudot, Ozan Ozisik

## Abstract

Environmental factors are external conditions that can affect the health of living organisms. For a number of rare genetic diseases, an interplay between genetic and environmental factors is known or suspected. However, the studies are limited by the scarcity of patients and the difficulties in gathering reliable exposure information.

In order to aid in fostering research between environmental factors and rare diseases, we propose ODAMNet, a Python package to investigate the possible relationships between chemicals, which are a subset of environmental factors, and rare diseases. ODAMNet offers three different and complementary bioinformatics approaches for the exploration of relationships: overlap analysis, active module identification and random walk with restart. ODAMNet allows systematic analysis of chemical - rare disease relationships and generation of hypotheses for further investigation of effect mechanisms.

**Metadata:** 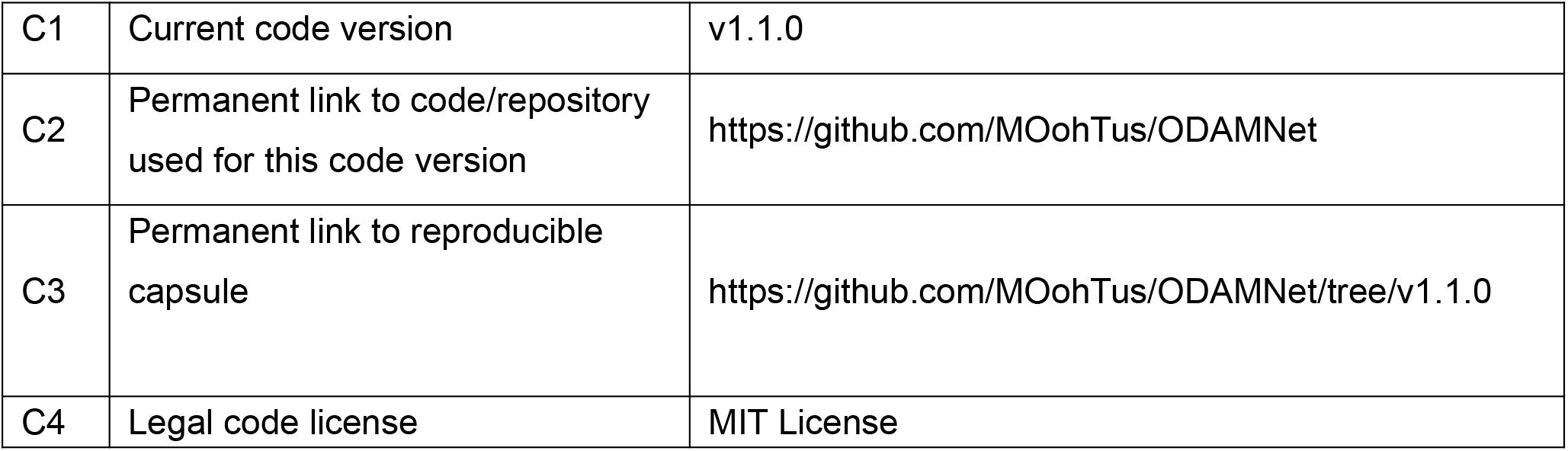

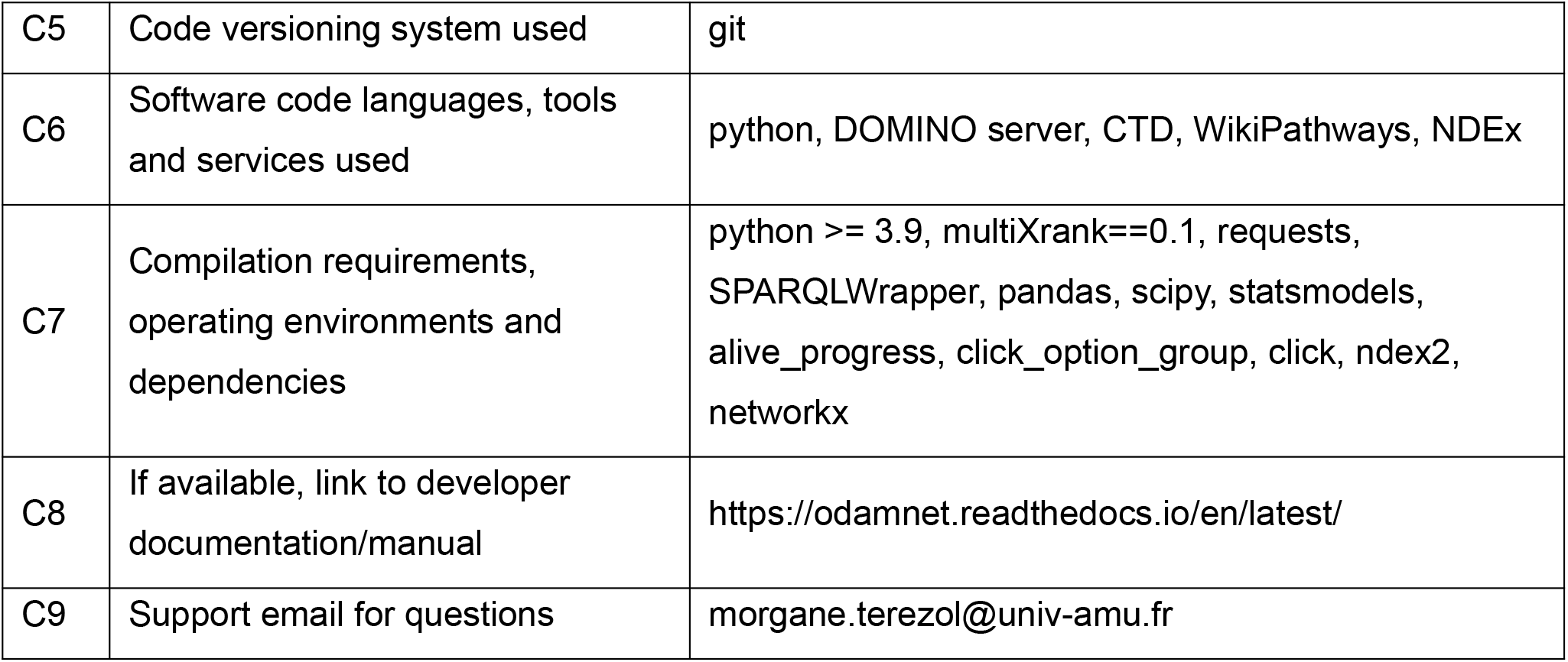

## 1. Motivation and significance

Environmental factors are external conditions that can affect the health of living organisms. These factors can be physical, biological, social, economic, or political [1]. They can play a significant role in the development and progression of genetic diseases. The relationship between genes and environment has been compared to the relationship between a loaded gun and its trigger [2].

The role of environmental factors have been reviewed for different diseases and disease groups, such as autism [3], inflammatory bowel disease [4, 5], cardiovascular disease [6, 7], congenital anomalies of the kidney and urinary tract [8, 9], amyotrophic lateral sclerosis [10], idiopathic pulmonary fibrosis [11], and Legg–Calvé–Perthes Disease [12].

There are multiple methodological challenges in studying the interplay between environmental factors and genetic diseases. Environmental factors have large spatial and temporal heterogeneities; a person’s activity patterns, residential changes and other conditions can modify the amount of exposure [13]. For the diseases that manifest later in life, relationships with an exposure that happened decades ago is hard to prove [10]. In the case of rare diseases, sample scarcity is adding another level of difficulty to the studies of environmental factors - disease relationships: low sample size and testing for multiple factors decrease the statistical power. Hence, limiting the environmental factors to be investigated is important, and this requires well supported hypotheses.

With this study, our aim is to provide a tool that can generate knowledge-based hypotheses regarding the relationships between chemicals, which are a subset of environmental factors, and the rare diseases. We previously investigated the role of vitamin A and vitamin D in the etiology of Congenital Anomalies of the Kidney and Urinary Tract (CAKUT) [14]. We explored the overlap between vitamin target genes and gene sets related to CAKUT. We observed significant enrichment of vitamin A target genes in CAKUT-related gene sets. Here, we propose ODAMNet (“Overlap, Diffusion, Active Module, Network”), a Python package that allows performing systematic analyses with i) multiple chemicals, ii) multiple rare diseases, and iii) multiple integrative bioinformatics approaches to explore the possible relationships between chemicals and rare diseases.

In ODAMNet, targets of the chemicals are retrieved from the Comparative Toxicogenomics Database (CTD) [15], and rare disease pathways are retrieved from WikiPathways [16]. We used three different and complementary bioinformatics approaches to integrate the data: overlap analysis, active module identification and random walk with restart. Of note, for the network-based approaches, i.e. active module identification and random walk with restart, the biological interaction networks are downloaded from the Network Data Exchange (NDEx) [17].

## 2. Software description

### 2.1. Software architecture

ODAMNet is a Python package for the investigation of chemical - rare disease relationships. It takes a list of chemicals as input and automatically retrieves the genes that are targeted by these chemicals from the Comparative Toxicogenomics Database (CTD) (Fig 1). Rare disease pathways are retrieved automatically from WikiPathways and networks are downloaded automatically from NDEx [17]. The user can also provide their own target genes, pathways of interest and biological networks. Then, ODAMNet can perform three different approaches. The first approach is an overlap analysis between the target genes and the rare disease pathways. The second approach is active module identification (AMI) using DOMINO [18], and then performs an overlap analysis between the identified active modules and rare disease pathways. The third approach is a random walk with restart (RWR) using multiXrank [19].

**Fig 1.**
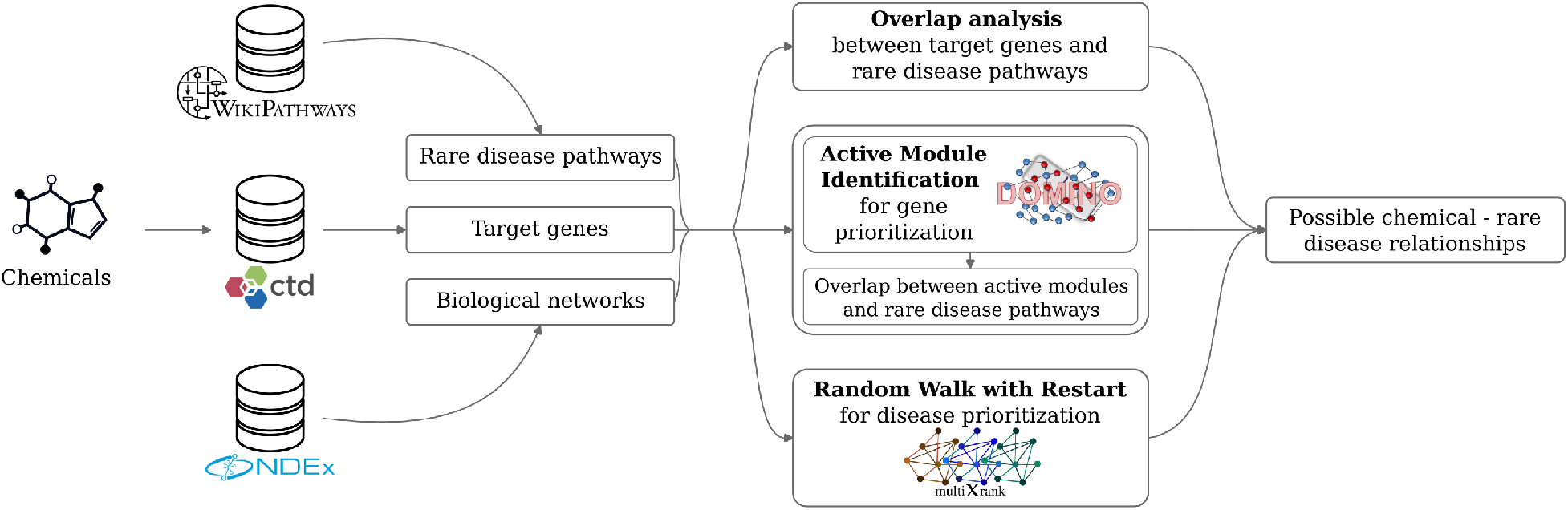
ODAMNet workflow

ODAMNet is written in Python 3 and is designed primarily as a command line program for the Linux operating system. It can be installed using pip and a detailed documentation is available on https://odamnet.readthedocs.io/en/latest/, with descriptions of:

- the three integrative bioinformatics approaches
- the format of input and output files
- all input parameters
- two use-cases, in which the datasets are either automatically retrieved or provided by the user

### 2.2. Software functionalities

#### 2.2.1. Data retrieval by queries

ODAMNet can retrieve the datasets required for an analysis, which are chemical target genes, rare disease pathways and biological networks, by querying the relevant databases. For the retrieval of genes targeted by a list of chemicals, the required input is a list of MeSH identifiers (https://meshb.nlm.nih.gov/) that correspond to those chemicals. Chemical target genes which have been reported for human are automatically retrieved from CTD (http://ctdbase.org/tools/batchQuery.go) using HTTP requests. The user can select the type of chemical -target gene associations. By default, *directAssociation* parameter is set to “True” and returns target genes solely for the input chemicals. Alternatively, setting this parameter to “False” will retrieve target genes not only for the input chemicals but also for any of its descendant chemicals. The user can also filter chemical - target gene associations based on the number of references, with the default being relationships supported by at least two references.

Rare disease pathways and the genes they involve are retrieved automatically from WikiPathways using SPARQL queries (https://sparql.wikipathways.org/). All human pathways and associated genes are also retrieved to construct the background gene set that will be used in the statistical tests.

In ODAMNet active module identification approach, biological networks are downloaded automatically from NDEx (https://www.ndexbio.org/) using the NDEx2 Python Client [20] and the networks’ universally unique identifiers (UUID). In the ODAMNet random walk with restart approach, multiple networks, including a bipartite network, are required. For this reason, ODAMNet does not perform the automatic download but provides assistive functions for downloading or building the necessary networks (please see section 2.2.4).

#### 2.2.2. Data input by the user

While the automatic retrieval of chemical targets and pathways from relevant databases is the default workflow, ODAMNet also allows the user to input custom target genes and gene sets such as annotation terms or pathways. In the case of using such custom gene sets, the user is expected to provide the background gene sets (as GMT files) that will be used in the statistical tests. It is possible to use gene sets from different databases (e.g. WikiPathways, Reactome [21], Gene Ontology (GO) [22, 23] in the same analysis; in this case, a background gene set for each database should also be provided.

As stated in the previous section, in random walk with restart, ODAMNet does not download and use networks automatically. However ODAMNet provides assistive functions to download networks from NDEx. The user can also provide their own networks. The user can provide their own network in the active module identification approach, as well.

#### 2.2.3. Integrative bioinformatics approaches

ODAMNet uses three approaches for finding the relationships between chemical target genes and rare disease pathways.

The first approach is an overlap analysis. It assesses if the chemical target genes are part of the rare disease pathways and applies a hypergeometric test to determine statistical significance. Benjamini-Hochberg method is then applied for the multiple testing correction.

The second approach is based on active module identification. An active module is a connected subset of gene nodes in a biological interaction network relevant to the investigated condition. For the discovery of active modules that have high connectivity and contain a high number of target genes (considered as active genes), we use the DOMINO method [18] with the default parameters through its web server [24]. Following the active module identification, a pairwise overlap analysis is performed between all the identified active modules and all the rare disease pathways. Duplicate significant rare disease pathways are removed, keeping only the most significant ones. The biological interaction network used by DOMINO can be automatically downloaded from NDEx or provided by the user.

The third approach is the random walk with restart (RWR) approach. RWR measures the proximity between given seed nodes (i.e., chemical target genes) and all the other nodes in the network, in a way analogous to a person that travels randomly on the connected nodes and sometimes teleports back to one of the seed nodes. RWR thereby considers both the network distance and the network topology. In the ODAMNet RWR approach, we add the rare disease pathways as nodes connected to their associated genes in the network. Then, RWR can find the rare disease pathway nodes that are proximal to the target genes, which are set as seeds. We use multiXrank v0.1 [19], a Python package that enables RWR on any kind of multilayer network. The biological networks are provided by the user. The input networks must include at least one network of genes/proteins, one rare disease pathways network (each node only connected to itself, disconnected otherwise) and a bipartite network connecting the rare disease pathway nodes to the genes in other networks. The rare disease pathways network and its corresponding bipartite network can be created using the *networkCreation* function available in ODAMNet. The user needs to provide a configuration file for multiXrank in which the seeds file and the network files are stated. multiXrank is run with the default parameters but these can be adjusted through the multiXrank configuration file. Please see our example configuration on https://github.com/MOohTus/ODAMNet/tree/v1.1.0, and multiXrank documentation at https://multixrank-doc.readthedocs.io and [19].

#### 2.2.4. Assistive functions

ODAMNet provides two assistive functions to download and build networks. The *networkDownloading* function allows downloading biological networks from NDex. This function uses the NDEx2 Python Client and the networks’ corresponding universally unique identifiers (UUID) provided by the user. The second function, *networkCreation*, allows the creation of the networks necessary for the ODAMNet RWR approach. It creates a rare disease pathways network (each node only connected to itself, disconnected otherwise) and its corresponding bipartite network which connects the rare disease pathway nodes to the genes.

## 3. Illustrative example

Vitamin A is a fat-soluble compound that plays an essential role in vision, intercellular communication, mucin production, embryogenesis, cell growth, and cell differentiation [25]. Deficiency or excess of vitamin A might play a role in disease development. Vitamin A deficiency can cause ocular degeneration, diverse changes in epithelial tissues, immune deficits, and excessive mortality from childhood diseases [25]. Excess or systemic intake of vitamin A during pregnancy can cause a spectrum of malformations including ocular, pulmonary, cardiovascular, and urogenital birth defects [25]. In this context, we used ODAMNet to investigate the molecular relationships between vitamin A and rare diseases. We used the automatic data retrieval functions of ODAMNet and its three analysis approaches.

### 3.1. Data query results

As input, we gave ODAMNet a chemicals file that contains the vitamin A MeSH ID (D014801). ODAMNet automatically retrieved the genes targeted by vitamin A and its 9 descendant molecules from CTD (*directAssociation* parameter set to “False”), which resulted in 7,765 human target genes for 10 chemicals. Then, we kept the chemical - gene associations with at least 2 references. After filtering, we obtained 2,143 target genes for vitamin A and its 6 descendants.

ODAMNet retrieved all pathways labeled as “rare disease” in WikiPathways, which resulted in 104 rare disease pathways. It also retrieved all human pathways to construct the background gene set. We obtained 1,281 human pathways in total.

The protein-protein interaction (PPI) network (UUID: bfac0486-cefe-11ed-a79c-005056ae23aa) was downloaded from NDEx. This network is the fusion of three datasets (Lit-BM, Hi-Union [26] and APID [27]). It is composed of 15,390 nodes and 131,087 edges.

The molecular complexes network (UUID: 419ae651-cf05-11ed-a79c-005056ae23aa) was also downloaded from NDEx. This network is the fusion of two molecular complex databases (CORUM [28] and HuMap [29]). It is composed of 8,497 nodes and 62,073 edges.

The last network downloaded from NDEx is the Reactome pathways network (UUID: b13e9620-cefd-11ed-a79c-005056ae23aa). This network was built based on data derived from Reactome protein-protein interaction data [21]. It is composed of 4,598 nodes and 19,292 edges.

All results are available on https://github.com/MOohTus/ODAMNet/tree/v1.1.0. Queries were made on 7 September 2022.

### 3.2. Overlap analysis results

We first used the ODAMNet overlap analysis approach. ODAMNet retrieved the chemical target genes and pathways as described in section 3.1. ODAMNet performed an overlap analysis between the 2,143 target genes and the 104 rare disease pathways. We obtained a significant overlap (adjusted p-value ≤ 0.05) between target genes and 28 rare disease pathways (Table 1).

**Table 1.**
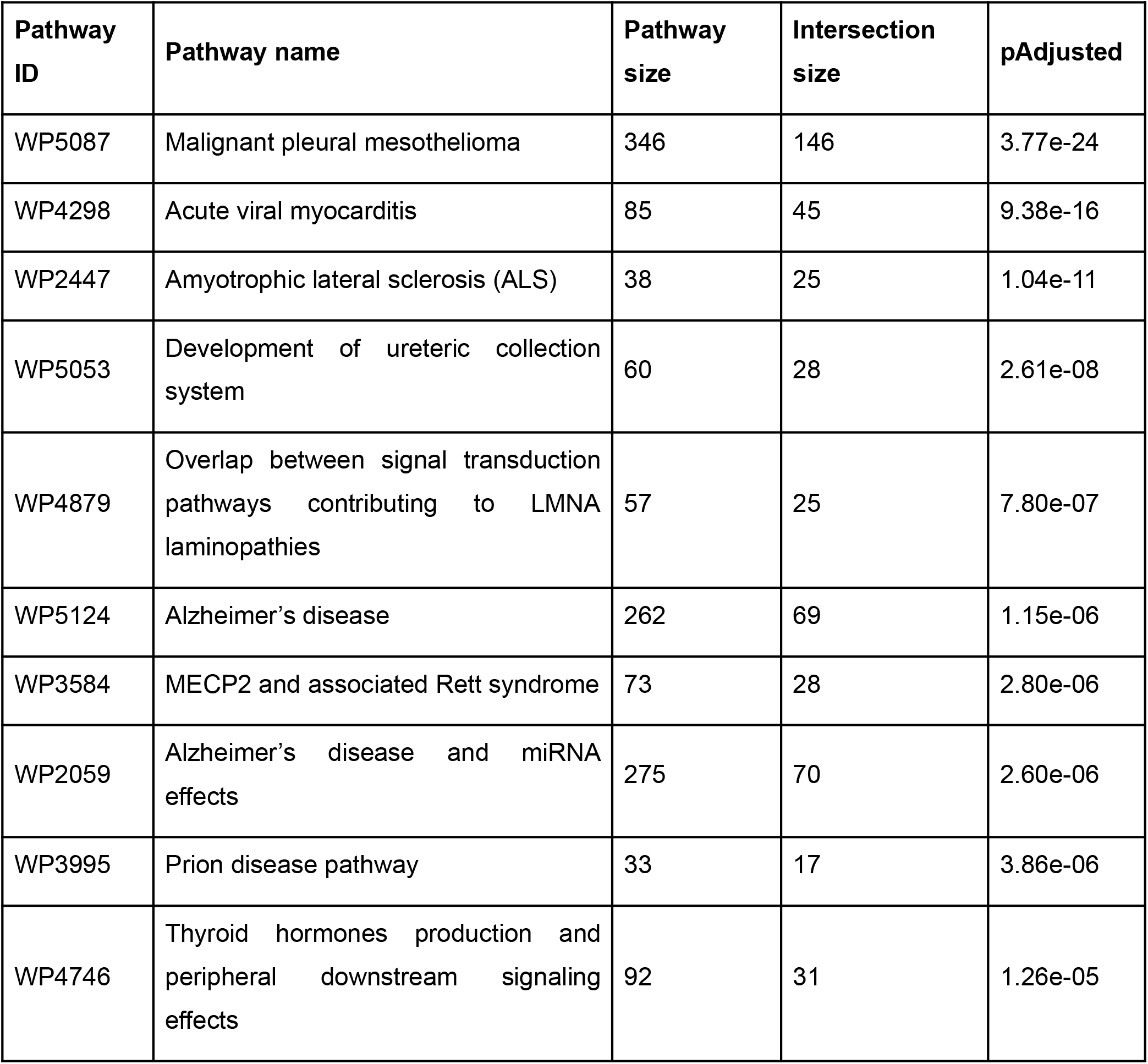
Top 10 of the 28 rare disease pathways significantly overlapping with vitamin A target genes.

The command used to perform this overlap analysis is:

~~~
odamnet overlap        --chemicalsFile useCases/InputData/chemicalsFile.csv \
                       --directAssociation FALSE \
                       --nbPub 2 \
                       --outputPath useCases/OutputResults_useCase1/
~~~

### 3.3. Active module identification results

To perform an active module identification approach, ODAMNet needs a chemicals file with the MeSH ID of vitamin A and the UUID of the PPI network. Then, for the overlap analysis between the identified active modules and rare disease pathways, ODAMNet extracted pathways from WikiPathways (see section 3.1 for more details about the query results). Over the 2,143 target genes, 1,937 were found in the PPI and used by DOMINO as active genes to find active modules. DOMINO found 12 active modules enriched in chemical target genes. Then, ODAMNet performed an overlap analysis between the 12 identified active modules and the 104 rare disease pathways. It found a significant overlap between 6 active modules and 19 rare disease pathways.

In Fig 2, we present 3 active modules as examples. We can observe that the topology of those active modules and the associated rare disease pathways vary. For instance, the active module on the right is highly connected and the genes are involved in many different rare disease pathways. The two other modules are less connected. The genes contained in the active module in the middle are involved only in the *Development of ureteric collection system* rare disease pathway.

**Fig 2.**
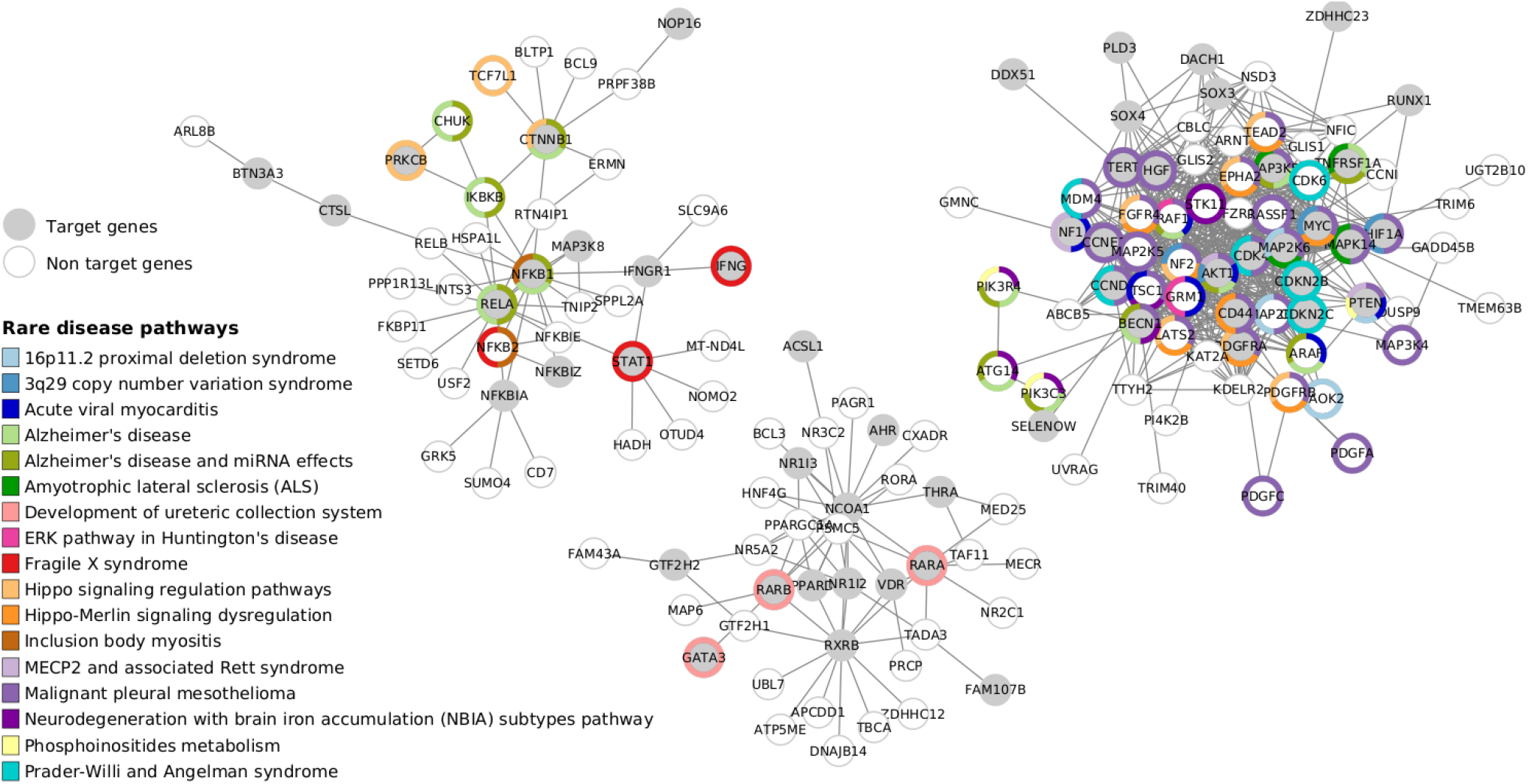
Active module identification with DOMINO followed by overlap analysis between genes in the module and the rare disease pathways. Visualization of 3 over 6 active modules identified by DOMINO. Target genes are in gray and non target genes are in white.

The command used to perform this active module identification is:

~~~
odamnet domino    --chemicalsFile useCases/InputData/chemicalsFile.csv \
                  --directAssociation FALSE \
                  --nbPub 2 \
                  --netUUID bfac0486-cefe-11ed-a79c-005056ae23aa \
                  --outputPath useCases/OutputResults_useCase1/
~~~

### 3.4. Random walk with restart results

To run the random walk with restart (RWR) analysis, we provided the vitamin A MeSH ID to automatically retrieve target genes from CTD (see section 3.1 for more details about the query results). We also provided a configuration file containing the path to the multilayer network. The multilayer network contains a gene multiplex network composed of three layers with the same nodes but different types of edges: the PPI network, the molecular complexes network and a network composed of Reactome pathways. We downloaded these networks from NDEx using the *networkDownloading* function. The multilayer network contains also a layer for the rare disease pathways network. This network layer contains disconnected rare disease pathway nodes (i.e., each node is only connected to itself), which are linked to the gene multiplex network by bipartite gene-disease associations. This rare disease pathways network and its corresponding bipartite network were created using the *networkCreation* function in ODAMNet.

MultiXrank used 2,012 chemical target genes available in the multiplex network (over the 2,143 genes retrieved from CTD). Then it calculated a RWR score for all the nodes of the multilayer network, which can be gene or rare disease pathway nodes. We selected the top 20 rare disease pathways nodes based on their RWR scores (Table 2).

**Table 2.** Top 10 rare disease pathways identified by the ODAMNet RWR approach for vitamin A

| Pathway ID | Pathway name | RWR score |
| --- | --- | --- |
| WP5087 | Malignant pleural mesothelioma | 2.85e-03 |
| WP4673 | Male infertility | 9.02e-04 |
| WP2059 | Alzheimer's disease and miRNA effects | 7.76e-04 |
| WP5124 | Alzheimer's disease | 7.76e-04 |
| WP4298 | Acute viral myocarditis | 7.69e-04 |
| WP3584 | MECP2 and associated Rett syndrome | 6.03e-04 |
| WP4746 | Thyroid hormones production and peripheral downstream signaling effects | 5.83e-04 |
| WP4549 | Fragile X syndrome | 5.77e-04 |
| WP5224 | 2q37 copy number variation syndrome | 5.66e-04 |
| WP4657 | 22q11.2 copy number variation syndrome | 5.62e-04 |

The command to download the PPI network used in the random walk with restart is:

~~~
odamnet networkDownloading        --netUUID bfac0486-cefe-11ed-a79c-005056ae23aa
                                  --networkFile PPI_network.gr
                                  --simple True
~~~

The command to create rare disease pathway network and its corresponding bipartite network is:

~~~
odamnet networkCreation        --networksPath useCases/InputData/multiplex/2/ \
                               --networksName WP_RareDiseasesNetwork_fromRequest.gr \
                               --bipartitePath useCases/InputData/bipartite/ \
                               --bipartiteName
~~~

Bipartite_WP_RareDiseases_geneSymbols_fromRequest.gr \

~~~
                           --outputPath useCases/OutputResults_useCase1
~~~

The command to perform this random walk with restart is:

~~~
odamnet multixrank     --chemicalsFile useCases/InputData/chemicalsFile.csv \
                       --directAssociation FALSE \
                       --nbPub 2 \
                       --configPath useCases/InputData/config_minimal_useCase1.yml \
                       --networksPath useCases/InputData/ \
                       --seedsFile useCases/InputData/seeds.txt \
                       --sifFileName resultsNetwork_useCase1.sif \
                       --top 10 \
                       --outputPath useCases/OutputResults_useCase1/
~~~

### 3.5. Comparison of the results obtained with the three integrative bioinformatics approaches

To summarize and compare the results from the three approaches, we used orsum [30], a Python package for filtering and integrating enrichment analysis results obtained from multiple studies (Fig 3). We observed that some rare disease pathways are identified as related to vitamin A by all three approaches, e.g. *Malignant pleural mesothelioma* and *Acute viral myocarditis*. Some others are specific to one or two approaches. For instance, *Male infertility* is retrieved with the overlap analysis and the random walk with restart analysis but not with the active module identification approach.

**Fig 3.**
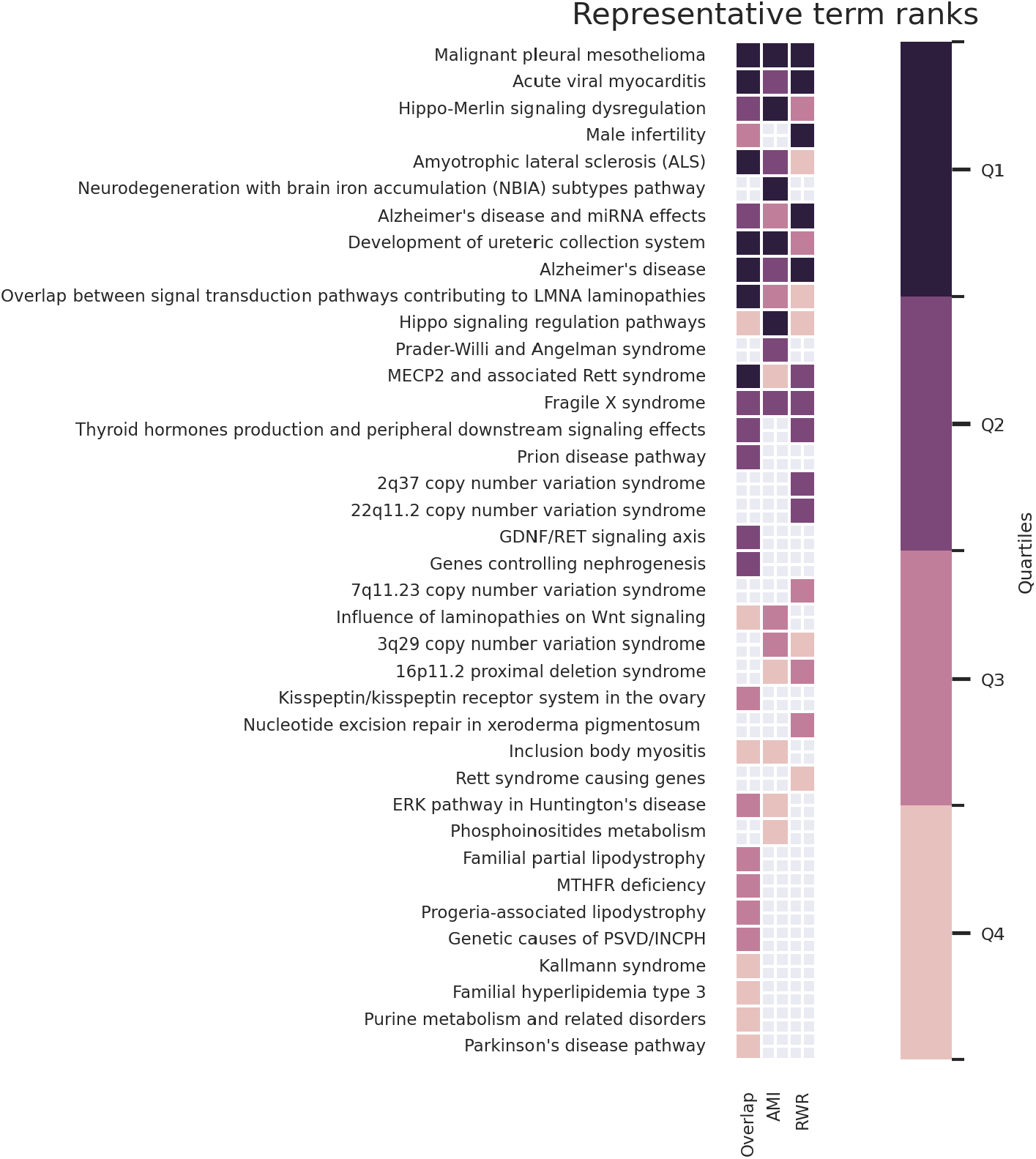
Summarization of rare disease pathways that are found to be related to vitamin A using overlap analysis (Overlap), active module identification (AMI) and random walk with restart (RWR) analysis. This heatmap is created by orsum [30].

## 4. Impact

ODAMNet offers a versatile and comprehensive framework for exploring the relationships between chemicals and rare diseases through its three distinct and complementary approaches. These three approaches work in different levels of knowledge: while overlap analysis focuses on the direct involvement of the chemical target genes in the rare disease pathways, active module identification considers both the target genes and the non-target genes in interaction with rare disease pathways. On the other hand, RWR finds the proximity of chemical target genes to the disease pathway nodes using more biological information through a multilayer network.

ODAMNet can be used in multiple ways to generate hypotheses for chemical - rare disease relationships. When investigating a specific chemical, this chemical can be tested for its relationships with multiple pathways, e.g. all the rare disease pathways from WikiPathways. Conversely, when investigating a specific disease, a set of chemicals and disease associated pathways can be determined and tests can be performed to assess their relationships. Furthermore, when the relationship between a chemical and a disease is already known, ODAMNet enables a focused analysis on the shared genes.

The automatic retrieval of chemical target genes, rare disease pathways and biological networks from relevant databases is the default workflow for ODAMNet. This ensures the usage of up-to-date data and enhances ease-of-use. ODAMNet also accepts direct custom input of target genes, gene sets and biological networks. This has multiple advantages. First of all, it allows reproducibility; target genes and rare disease pathways from specific versions of CTD and WikiPathways can be stored and used in ODAMNet. Second, even if the external resources become temporarily unavailable, ODAMNet can still be run with the previously saved data. And last, this functionality extends the possible use-cases of ODAMNet, as any gene list and gene set data can be incorporated.

## 5. Conclusions

Environmental factors, in particular chemical substances, are known or suspected to be playing a role in multiple diseases. There is growing knowledge on chemical - gene/protein interactions and rare disease molecular mechanisms. We designed ODAMNet to bridge the gap between these knowledge sources. Using three different and complementary approaches, ODAMNet can generate knowledge-based hypotheses for chemical - rare disease relationships and their underlying mechanisms. Overall, ODAMNet’s comprehensive approach, flexibility in analysis design, and utilization of up-to-date or custom input data should allow its usage in a wide variety of contexts.

## Acknowledgements

We thank the members of the European Joint Programme on Rare Diseases for their support and discussions. We thank Nazli Sila Kara for being one of the first users of ODAMNet and giving much useful feedback. We thank Cécile Beust who created the biological networks and uploaded them to NDEx.

## Author contributions

Morgane Térézol: Methodology, Software, Writing - Original Draft, Writing - Review & Editing, Visualization.

Anaïs Baudot: Methodology, Writing - Review & Editing, Supervision, Project administration.

Ozan Ozisik: Methodology, Writing - Original Draft, Writing - Review & Editing, Supervision, Project administration.

